# Spatial Patterns for Discriminative Estimation

**DOI:** 10.1101/746891

**Authors:** Alberto Llera, Roselyne Chauvin, Peter Mulders, Jilly Naaijen, Maarten Mennes, Christian F. Beckmann

**Author notes:** AL,; RC,; PM,; JN,; MM,; CB,.

## Abstract

Functional connectivity between brain regions is modulated by cognitive states or experimental conditions. A multivariate methodology that can capture fMRI connectivity maps in light of different experimental conditions would be of primary importance to learn about the specific roles of the different brain areas involved in the observed connectivity variations. Here we detail, adapt, optimize and evaluate a supervised dimensionality reduction model to fMRI timeseries. We demonstrate the strength of such an approach for fMRI data using data from the Human Connectome Project to show that the model provides close to perfect discrimination between different fMRI tasks at low dimensionality. The straightforward interpretability and relevance of the model results is demonstrated by the obtained linear filters relating to anatomical areas well known to be involved on each considered task, and its robustness by testing discriminatory generalization and spatial reproducibility with respect to the number of subjects and fMRI time-points acquired. We additionally suggest how such approach can provide a complementary view to traditional task fMRI analyses by looking at changes in the covariance structure as a substitute to changes in the mean signal. We conclude that the presented methodology provides a robust tool to investigate brain connectivity alterations across induced cognitive changes and has the potential to be used in pathological or pharmacological cohort studies. A publicly available toolbox is provided to facilitate the end use and further development of this methodology to extract Spatial Patterns for Discriminative Estimation (SP♠DE).

## I. INTRODUCTION

Functional magnetic resonance imaging (fMRI) is the primary tool used to investigate how the brain (re)acts under different mental states and the biological underpinnings of brain disorders. Most commonly, these state or group-differences are investigated using generalized linear models (GLM) [1] to detect brain regions that differentiate between experimental conditions. The GLM is for example the standard tool used to analyze task fMRI data incorporating a temporal model for the induced brain activity [2] that allows obtaining a supervised voxel-wise characterization of changes in the blood-oxygen-level dependent (BOLD) response [1]. Yet, while powerful and straightforward to interpret, the GLM has two crucial disadvantages. Foremost, while fMRI data is multivariate in nature, the GLM is applied as a mass univariate approach (i.e., independently to each voxel in the brain) and does not consider interactions between different regional effects. Second, it ignores the variance in the data by looking at single mean BOLD changes across conditions. To overcome these issues, one can use multivariate models that allow studying the second order interactions between different brain areas [3], [4]. Although these models overcome some of the issues with univariate modelling, they are not specifically tailored for state/group characterization. Further inclusion into models for state/group identification, i.e. regression or classification, can provide strong discrimination but complicates the interpretation of the results. Consequently, to improve interpretability of results obtained from analyzing multivariate fMRI data while optimizing discrimination between two conditions or groups of fMRI time-series, it would be beneficial to the community the use of multivariate approaches to extract brain patterns that maximize discriminative properties. Using a simultaneous diagonalization approach [5] we introduced such preliminary results on fMRI data in [6] and further work has been done since then by applying it to cohorts of neurodevelopmental disorders [7]–[9]. Recently we have also considered it to study the effect of stress induction [10]. For a proper understanding between the fMRI community, we first summarize here the full analytical transformations defining the simultaneous covariance diagonalization [5]. The basics of the general approach [5] are not new to the neural engineering community as it is commonly used as a feature extractor for classification on EEG-based brain-computer interfaces [11], [12]. In such prediction contexts, the model order selection, i.e. which subset of linear filters to select, represents no critical choice due to the posterior features interaction with a classifier, and simple heuristics commonly suffice. However, when considering such model further than for discrimination, the clinician/researcher needs to know which set of filters deserve further attention. In addition, the model order selection step is required for the interpretation of spatial maps derived from ‘backward’ linear models [13], and consequently, using simultaneous diagonalization approaches further than for discrimination, but as a cognitive neuroscience tool, requires stricter model order optimization. Here we embed the model order optimization into a permutation testing statistical approach. However, the application of this model to fMRI data is principally challenging due to the high spatial correlation of fMRI data and because the model requires a large number of observations to compute well-posed covariance matrices. Although recent advances in data acquisition and large-scale data collection allow collecting a large number of observations in relatively short time periods, this is not yet the standard, especially in clinical acquisitions. To partially overcome these issues, here we perform an initial fMRI spatial dimensionality reduction using a functional brain parcellation [14], and compute regularized versions of the spatial covariance matrices [15] to expand the original methodology formulation [5]. We make use of large-scale high-quality fMRI data from the Human Connectome Project (HCP) [16], [17] to uncover and evaluate the potential power of such pipeline for discriminative estimation and spatial reproducibility on fMRI data. We consider resting state, motor, and working memory task fMRI data from 1063 subjects and demonstrate that we are able to learn robust low-dimensional representations of the data that provide close to perfect discrimination between the different mental states. The method is fast and efficient and the availability of these simple and well-known tasks allows to validate that the learned filters are robust and specific to brain areas well-known to be involved during their respective paradigms [13], [18], validating the spatial interpretation of the obtained results on fMRI data. Further, the highly temporally sampled fMRI data of the HCP allows also considering the discrimination and spatial associations’ robustness as a function of the number of subjects and time points included. Herewith, identifying the scenarios in which the current approach is useful to address general questions using fMRI.

## II. Methods

### A. Dataset & preprocessing

In this work we use resting state and task fMRI data from the Human Connectome Project (HCP) [16], [17]. For each subject, we consider the motor task (MT), the working memory task (WM), and the first of the two available resting state sessions (RS). We use the HCP1200 release which includes data from 1200 subjects and only consider data from subjects for whom all three data modalities (RS, MT and WM) were available resulting in a total of 1063 subjects. In all cases we used the fully preprocessed data in MNI152 space as delivered in the HCP1200 release. For the full preprocessing details we refer the reader to [19].

### B. Spatial Patterns for Discriminative Estimation

Next, we describe the processing steps that lead to the extraction of Spatial Patterns for Discriminative Estimation (SPADE) from fMRI data. SPADE indexes connectivity changes from two sets of fMRI data using the simultaneous diagonalization of two covariance matrices [5]; full analytical details are provided in Appendix A. Although ideally one would like to perform such analyses using well posed full brain spatial covariance matrices, this is not achievable yet due to the fMRI spatial correlation and the computational memory constraints relating to the high spatial resolution of fMRI measurements. As a consequence, a spatial dimensionality reduction must be performed in order to apply the algorithm to fMRI data. For each subject and for each of the three considered fMRI data modalities we performed a spatial dimensionality reduction into 165 regions of interest (ROI) from a functional parcellation [14] by extracting the mean time-series across the voxels in each ROI. Each ROI time-series was then independently demeaned and divided by its standard deviation before further processing.

Then, considering such ROI time-series gathered under two different fMRI modalities (e.g., resting state versus working memory task fMRI), we compute a regularized covariance matrix[15] per subject and condition and average then across subjects to obtain a unique covariance matrix per modality; these two average covariance matrices are then used to estimate their simultaneous diagonalization[5]. This results in a set of discriminative linear spatial filters that optimally separate the two initial modalities in terms of variance. We address the model order selection, i.e. the number of spatial filters selected for discrimination, using permutation testing. Briefly, we select the filters for which the difference of the variances across tasks rejects the hypothesis of their variances being equal for both tasks. This is achieved by building a null distribution by balanced permutating the data across conditions, and we used 1000 permutations (see Appendix C). An important feature of the SPADE model is that connectivity changes can be summarized for each basis vector as a unique spatial map that provides a spatial weight for each ROI. This results in a straightforward interpretation of the relevant changes in connectivity. Since the simultaneous diagonalization assumes no implicit noise model, the estimated basis vectors cannot be directly interpreted back to the brain, and the interpretable associated spatial maps are obtained using structural coefficients [13], [18]. Brief details are provided in Appendix B and for a more extensive explanation we refer the reader to [13]. Note that since for a proper spatial interpretation the source model is estimated from the selected relevant filters [13], the model order selection we introduce here is of relevance to obtain a proper interpretation of the resulting spatial filters.

For the remainder of the paper we will denote the full process involving spatial dimensionality reduction of two fMRI data modalities, spatial covariance regularization, simultaneous diagonalization, model order estimation by permutation testing, and estimation of the associated interpretable spatial maps [13] as SPADE. To facilitate the end use application of these tools to the community, a toolbox providing full automatized estimation will be made publicly available through Git-Hub upon publication.

### C. Evaluations

We use the resting state (RS), motor task (MT) and working memory task (WM) fMRI data from the HCP sample and apply the introduced SPADE methodology to find brain connectivity differences between each pair of fMRI modalities independently: RS vs MT, RS vs WM and MT vs WM. To quantify the quality of the learned filters we perform a discriminative analysis to distinguish between fMRI modalities using a ten-fold cross validation approach for each of the three scenarios (RS vs MT, RS vs WM and MT vs WM) independently. At each fold, SPADE discriminative filters are learned from the covariance matrices of the two selected modalities using data from 90% of the subjects and the number of filters is selected using permutation testing. Then we compute the projection of these subjects’ data into the newly learned basis, compute the logarithmic variance of the resulting time-series as features, and use them to train a Linear Discriminant Analysis (LDA) classifier [20] to distinguish between the two included modalities (e.g. RS vs MT). The filters are then used to project the remaining unseen data (~10% subjects) and extract the log-variance of these projections as features to test the LDA classifier quality. Each classifier is evaluated using the accuracy, i.e. the percentage of correctly classified samples, and we report statistics across folds. We then test the significance of mean classification performance against a null distribution built from accuracy values obtained from SPADE analyses where the labels where evenly mixed across samples (1000 randomizations). For comparison we also evaluate the results obtained using a linear support vector machine (SVM) [21] classifier in the full covariance space, i.e. feature space of dimension (165*166)/2= 13695. To conclude the analyses of these full samples we visually evaluate the spatial extent of the SPADE linear filters using structural coefficients for visualization and interpretation of the associated spatial maps [13], [18].

Then, we consider the effect of a reduced number of subjects and temporal observations in the performance of the model. For each considered number of subjects (N) and time-points (T), we performed 1000 different bootstraps using the SPADE analyses, where at each realization a subset of N subjects was selected randomly and the temporal down-sampling was performed by selecting T equally temporally spaced samples from the full original fMRI time-series i.e. simulating higher TR acquisitions. The performance of the model at these scenarios was assessed using classification performance and spatial reproducibility. To evaluate the spatial reproducibility of the results we first compute the correlation between the spatial maps obtained in the original decomposition and the ones obtained at each bootstrap. We then estimate the probability of each map being significantly recovered by computing the percentage of bootstraps for which the correlation value rejects the hypothesis of not being correlated at a given significance level i.e. testing against a null distribution build from the off-diagonal terms of the correlation matrices across bootstraps.

In addition, we perform a secondary evaluation where we apply the SPADE model to a scenario previously studied using a GLM analysis. Using the WM task data from the HCP sample, we compared the 0-back memory task periods to the 2-back memory task periods using the SPADE approach. As with the previous evaluations, in this case we also test the discrimination power using 10-fold cross-validation and report the group spatial maps associated to the top four spatial maps for comparison to the results presented in [22].

## III. Results

To illustrate the discriminative performance of the SPADE methodology, Fig. 1 presents two-dimensional representations of fMRI data obtained using the SPADE model.

**Fig. 1:**
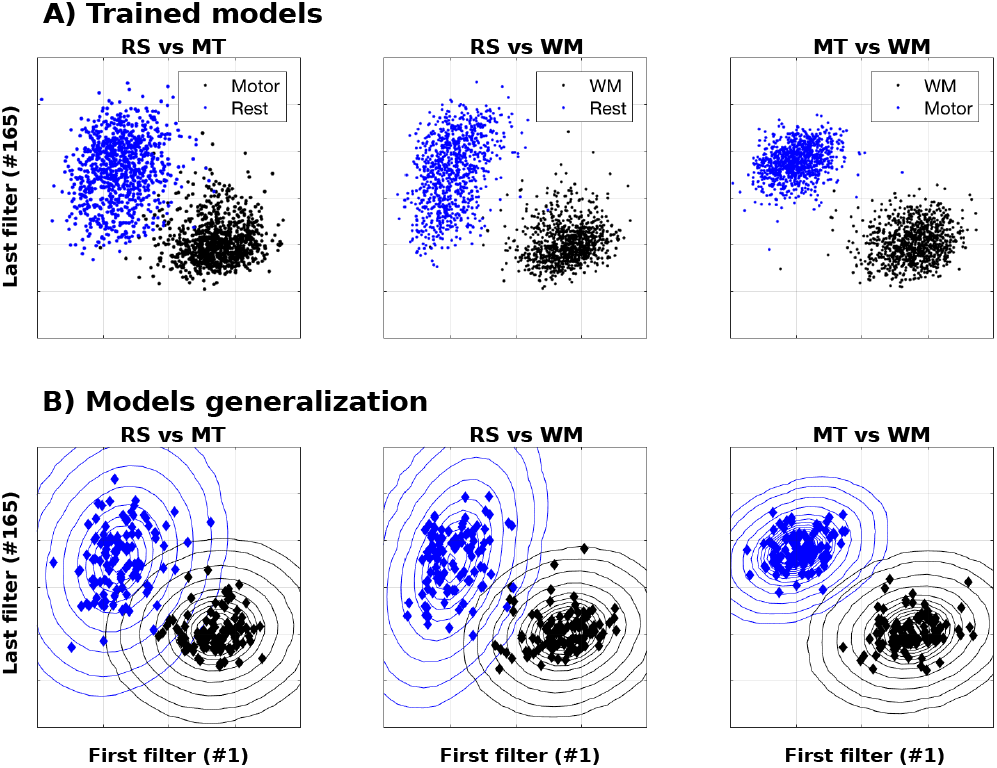
Two-dimensional fMRI data representations obtained using SPADE. The columns represent three different comparisons, MT vs RS, WM vs RS and WM vs MT task fMRI respectively. In the first row we present the logarithmic variance of the training fMRI data projected onto the first and last SPADE filters learned to discriminate each pair of fMRI modalities as indicated in the figure legends. Dots represent subjects and blue and black color encode the two different fMRI tasks considered on each scenario. In all three cases we can see that the training set is correctly separated in two visually distinguishable sets. In the second row we represent the learned two-dimensional training feature distributions presented in the first row as two Gaussian densities, and visualize the logarithmic variance of the testing fMRI data projections into the new basis as color coded diamonds. Appreciate the generalization ability of the SPADE model that provides an optimal twodimensional classification space.

For each comparison between two fMRI modalities, we visualized one random fold from the ten-fold-cross-validation used to evaluate discrimination and divide the data into training and testing datasets. Note that by construction, the first and the last SPADE filters maximize variance for one class while minimizing it for the other one. Consequently, the discriminative power of the algorithm usually comes from the combination of pairs of filters at both extremes of the eigen-spectrum. In the first row we can appreciate, for each of the three comparisons, a very clear two-dimensional separation between the training features obtained by combining the two extreme SPADE filters. The second row clearly shows the generalization ability of the SPADE model on the test-set data. We observed that in all cases the classification using uniquely one pair of filters (two-dimensional feature space) is very high, with a mean value above 99%; exact statistics are provided in the second column of Table 1.

**Table 1:** Summary results of the SPADE analyses. The second row presents the classification accuracy results when comparing resting state (RT) with motor task (MT) fMRI, the third row shows RT with working memory (WM) task fMRI, and the forth row compares WM and MT fMRI. The fifth row shows the mean results across the three comparisons. The second column (SPADE 2-D) presents the classification results (mean and standard deviation) when discriminating each pair of fMRI modalities using a SPADE two-dimensional projection of the data. The column ‘SPADE selected filters’ shows the eigen-spectrum location of the filters selected by the model order selection. The column ‘SPADE accuracy’ shows the classification results obtained using the model order selection and the column ‘SVM accuracy’ shows the results obtained using a SVM in full covariance space (i.e. feature space of dimension (165*166)/2= 13695).

|  | SPADE<br>2D<br>accuracy | SPADE<br>Selected<br>filters | SPADE<br>accuracy | SVM<br>accuracy |
| --- | --- | --- | --- | --- |
| RT vs MT | 98.64%<br>(0.95) % | 1-6<br>155-165 | 99.3%<br>($2 \cdot 10^{-3}$) % | 98.9%<br>(0.37) % |
| RT vs WM | 99.34%<br>(0.33) % | 1-2<br>152-165 | 99.77%<br>($6 \cdot 10^{-5}$) % | 99.8%<br>(0.40) % |
| WM vs MT | 99.76%<br>(0.46) % | 1-4<br>153-165 | 99.63%<br>($2 \cdot 10^{-4}$) % | 99.58%<br>(0.35) % |
| mean | 99.24%<br>(0.58) % | | 99.56%<br>($1.3 \cdot 10^{-3}$) % | 99.42%<br>(0.37) % |

We then applied the proposed model order selection via permutation testing and selected filters at significance Bonferroni corrected level (p<0.05/165 = 3.3×10^-4, see Appendix C). Although in practice the dimensionality was computed on each fold independently, for illustration we provide the exact set of filters selected when using the full sample at each comparison in Table 1 column ‘SPADE selected filters’. On each of the three comparisons the classification performance obtained using SPADE at the dimensionality selected using the permutation testing strategy is presented in Table 1 column ‘SPADE accuracy’. In all cases, SPADE classification performance was significantly better than random (permutation p<10^-3^). For comparison, we also provide the classification results obtained using an SVM in the full covariance space in the last column of Table 1. Although the mean improvement with respect to the two-dimensional scenario or the SVM is relatively small, the advantage of using model order selection for discrimination is most notable in the reduced standard deviation of the results across folds. These results highlight that the top filters are the most relevant for discrimination but using the model order selection generalizes better across folds. Further, the comparison with the SVM provides remarkable results, especially in light of the reduced dimension in which SPADE classification is performed and the linear nature of the SPADE filters that allow for a straightforward spatial interpretation.

To evaluate the spatial distributions of the filters associated to the discriminative results reported, Fig. 2 summarizes the spatial maps obtained in all three comparisons. Since each fold provides slightly different discriminative filters, for visual validation we performed SPADE analyses including all subjects in the learning phase. The obtained spatial filters were then transformed into interpretable spatial maps [13], [18].

**Fig. 2:**
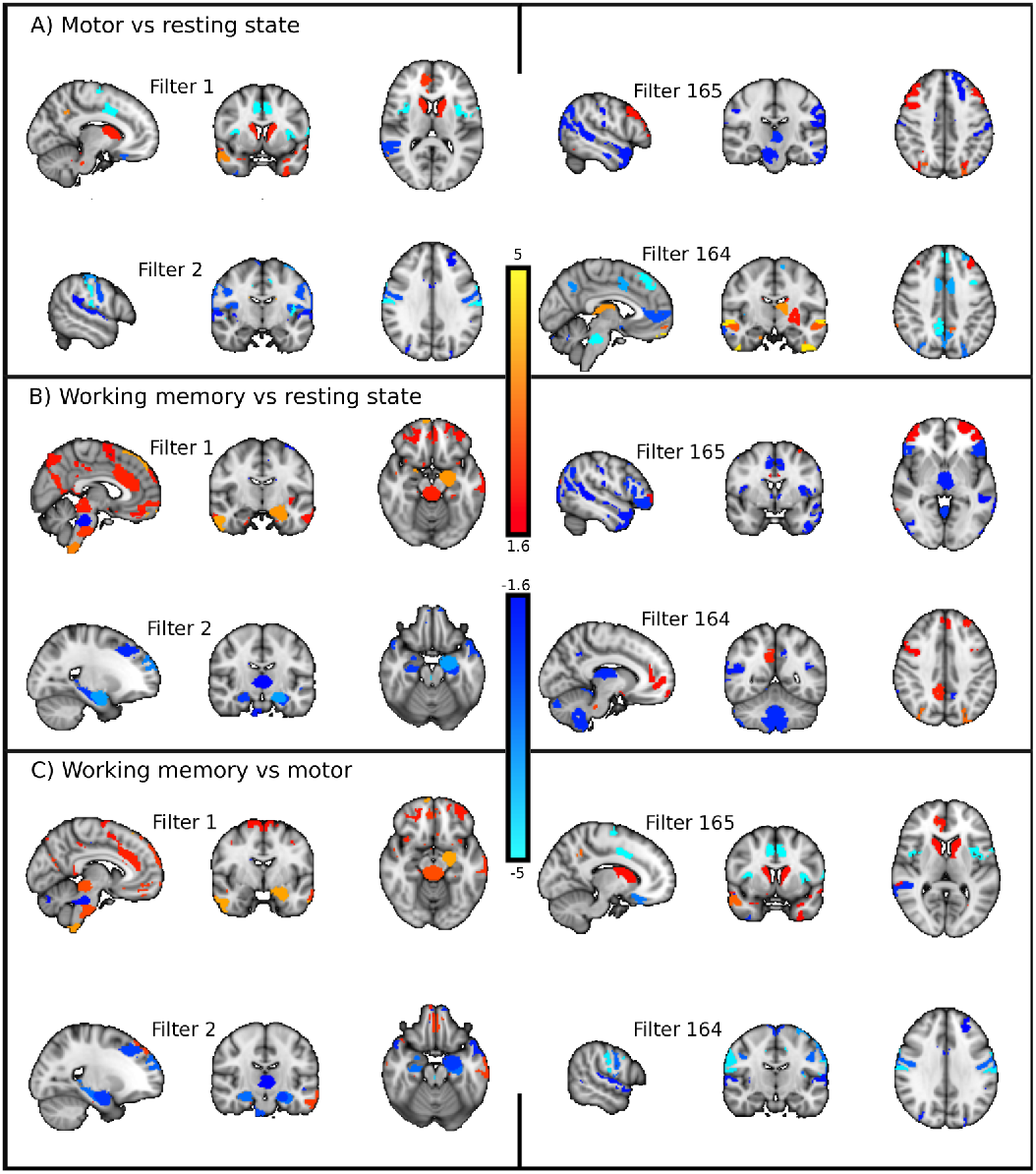
A) Resting state vs. motor task. B) Resting state vs. working memory task. C) Working memory vs. motor task fMRI. For each of the three scenarios we present a summary of the spatial maps associated with the discriminative spatial filters. On the left side we present filters 1 and 2, which maximize variance for A) motor, B) memory and C) memory. On the right side we present filters 165 and 164, which maximize variance for A) resting state, B) resting state and C) motor task. For visualization the value of the maps is converted to pseudo-z-statistics and thresholded at Z=1.6.

When looking at differences between resting state and motor task [23], in Fig. 2A) we observe that areas synchronizing to maximize variance for the motor task (left side, filters 1 and 2) involve bilateral supplementary motor cortex, caudate and operculum in the first filter, while primary motor cortex and contralateral cerebellar cortex involvement is present in the second filter. Areas identified by these filters are well known to be related to motor preparation, execution and control [24], [25]. On the other side of the spectrum (right side, filters 164 and 165) we find the filters maximizing variance for resting state data while minimizing it for motor task data. The last filter (#165) involves sensory motor cortex, bilateral dorso-lateral PFC (attention network), cerebellum and several temporal areas including the temporal gyrus and temporal pole. On the other side, the penultimate filter (#164) resembles the default mode network (DMN), anterior cingulate cortex and thalamus.

When looking at the differences between resting state and working memory task, in Fig. 2B) we observe that the areas synchronizing to maximize variance for the working memory task in the first filter involve areas well known to working memory performance [26], [27] as visual and precuneus area, similar to the dorsal pathway, and temporal-occipital areas, as present in the ventral pathway [28], [29]. Filter number two involves areas similar to number one but includes bilateral hippocampus and thalamus and more temporal areas. On the other side of the eigen-spectrum, we find that the last filter (#165) involves the anterior cingulate cortex, cerebellum, frontal pole, thalamus, middle temporal gyrus and temporal pole, frontal opercula. Further, the penultimate filter (#164) involves again the DMN and the thalamus. We observe that the filters relating to increased variance during rest (Fig. 2A and 2B) are in both cases informing about a stronger involvement of default mode network (DMN) related areas during the resting state and are extremely similar in both independent analyses (Fig. 2A and B right sides), showing the consistency of the SPADE algorithm.

When looking at the differences between working memory and motor task (Fig. 2C), we observe that the spatial maps obtained are also very closely related to the ones that differentiated the tasks from the resting state; in particular, filters maximizing for WM (Fig. 2C, filters 1 and 2) resemble the ones maximizing variance for WM with respect to resting state (Fig. 2B filters 1 and 2); and filters maximizing variance for MT (Fig. 2C, filters 164 and 165) exhibit a similar spatial distribution to the ones maximizing variance for MT with respect to resting state (Fig. 2A filters 1 and 2). These results show that the filters extracted for each modality involve areas well known to be required for the particular task performed and are highly specific since they can be obtained independently of the other covariance structure considered.

We then consider the behavior of the model as a function of the number of subjects and of fMRI time points. For each of the three considered comparisons, Fig. 3 presents the mean and standard deviation (top and bottom rows respectively) of the classification performance results as a function of the number of subjects and of fMRI time points included, (see section Methods for details). In general, all the classification performances above 60 % where found significant at the granularity level (permutation p<10^-3^). Clearly more subjects and longer scanning sessions provide stronger discriminability as shown by a higher accuracy and a lower standard deviation in these numerical experiments. However, we appreciate that in all cases above 100 subjects and a reasonable amount of time points (350 as a reference) are enough to reach close to optimal performance and a low standard deviation. Using 50 subjects still provides high accuracy in most cases but at a very high standard deviation cost. Dropping the number of subjects to 20 or less clearly decreases the model performance even when considering a high temporal sampling.

**Fig. 3:**
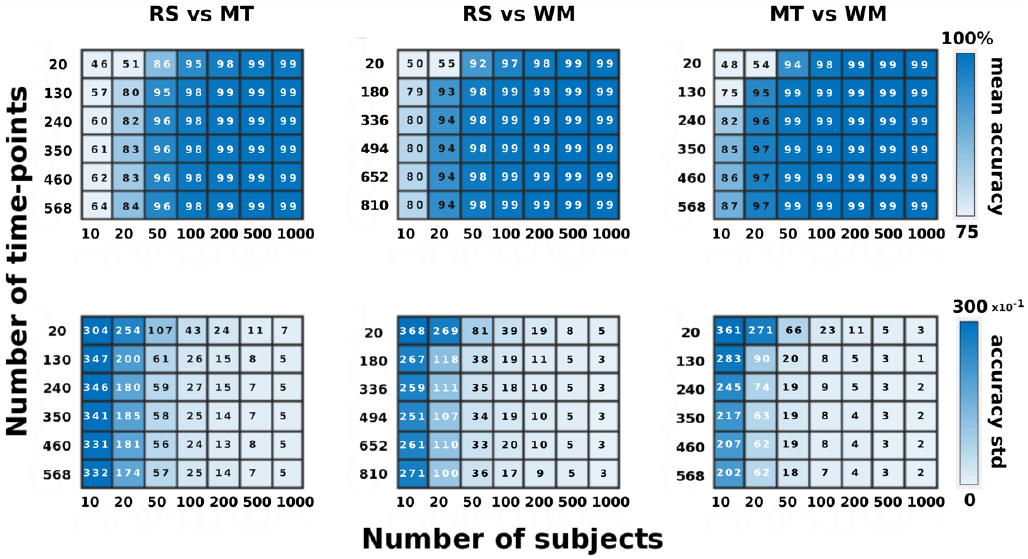
Top row presents the mean classification performance for the three comparisons indicated on each figure title, each as a function of the number of fMRI time samples (y-axis) and the number of subjects (x-axis). The bottom row shows the standard deviation around the classification accuracy presented in the top row.

The numerical results presented in Fig. 3 are of interest when the goal of a study is to discriminate between two groups. However, our main aim is to develop models to learn about the hidden brain sources that drive observed changes on brain functional connectivity as a function of cognitive changes, and consequently, we are required to address the reproducibility of the spatial maps obtained through SPADE. For space constrains we present detailed results only for the RS vs MT case; the results for the other two scenarios, RS vs WM and MT vs WM were quantitatively but not qualitatively different. Fig. 4 shows the mean correlation matrices between the significant spatial maps obtained in the full sample (N=1063) and the spatial maps obtained for the number of subjects indicated on each subfigure title (N) and the number of time-points indicated in the y-axis (T) (see Methods). The standard deviations around these mean values are shown in Appendix D, Fig. A1. The size of these matrices equals the number of significant selected filters indicated in Table 1 since all the analyses were performed at the same dimensionality than the full analyses (see Table 1). Further, the obtained maps at each subsample were not reordered to match the original order.

**Fig. 4:**
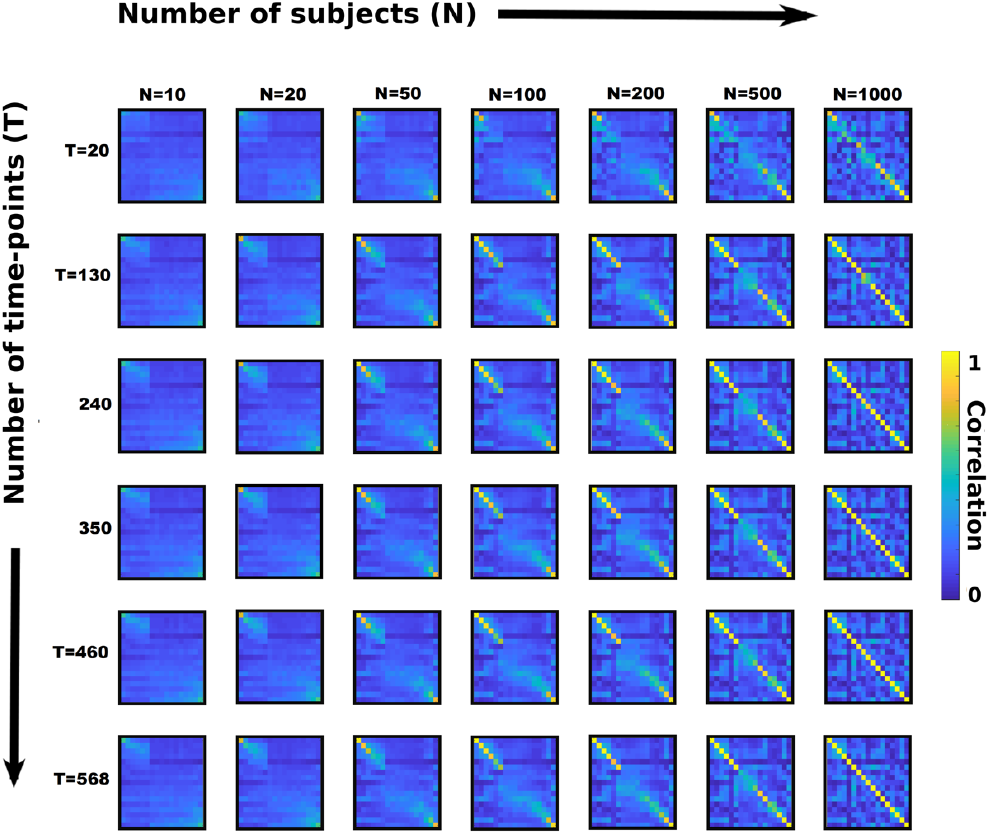
SPADE spatial maps reproducibility for RT vs MT. Each subfigure shows the mean correlation matrix between the significant spatial maps using the full sample (x-axis, N=1063) and the ones obtained for N and T as indicated on each row and column (y-axis). The mean is computed over 100 random selections of subjects (see Methods section for details). The size of each correlation matrix equals the number of significant selected filters as indicated in Table 1 and, are accordingly ordered. In this way, the top left and bottom right corners of each subfigure relate to the spatial reproducibility of the most important filters, the first (#1) and the last one (#165) respectively.

For each subfigure, an identity matrix would represent perfect reproduction of all selected spatial maps and consequently, it becomes clear that as with the classification performance, the reproducibility also increases stronger with higher number of subjects than with longer scanning sessions. The top left and bottom right corners of each subfigure relate to the spatial reproducibility of the two most discriminative spatial maps, the associated to the first and the last filters. We observe that these two spatial maps start to be reproduced relatively well in expectation with N=50 subjects at relatively short scanning sessions, and that in general it seems to be easier to recover maps in the top of the eigen-spectrum. However, since it is difficult to draw conclusions from such representation, in Fig. 5 we show for each N and T, the probability of each spatial map being successfully reproduced (y-axis), at different significance levels (x-axis) (see Methods for details). For visualization the results are shown explicitly for the first, second, penultimate and last spatial maps (1, 2, 164, 165), and for all other spatial maps/filters we show the mean and standard deviation of a) all other significant maps (green coded) and b) all non-significant maps (orange coded).

**Fig. 5:**
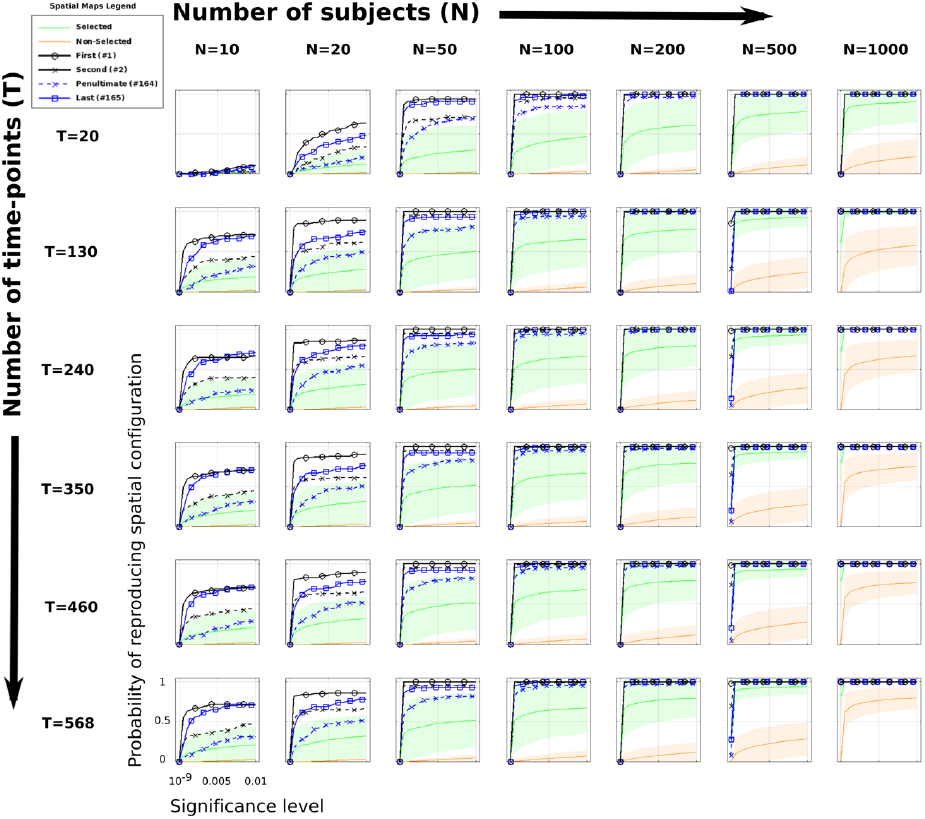
Probability of obtained spatial maps being significantly reproduced (y-axis) at a given significance level (x-axis), for each N and T. Black and blue lines encode the top four strongest filters as indicated in the figure legend. Colored areas encode the mean and standard deviation obtained across the significantly selected maps (green) and the non-selected filters (orange).

This representation shows that we are able to recover the first and last filters with 50 subjects, but also that the last filter is not reproduced with probability 1, even at (uncorrected) p < 0.01. Further, we observe that reproducing the penultimate filter required at least 100 subjects. When considering the reproducibility of the other significant maps (green coded), we see that an accurate reproduction is achieved at N=500. It is to note that at N=500, and even at N=1000, the remaining nonsignificant filters (orange coded) are poorly reproduced, reflecting the heterogeneity of the non-tasks discriminative data sources across subjects, and remarking the reproducibility of the task related selected components. Altogether, these results show that the spatial reproducibility of the results obtained when using this model is dependent on the set of filters to report. As a reference, when requiring simply the two extremer maps (first and last), a set of N=50 subjects would be sufficient. However, we would need around 100 subjects to interpret or include in further analyses the four strongest maps, and around 500 subjects to recover fully the set of significant spatial maps. Further, although there is a preference for longer scanning sessions, scans around 350 timepoints seemed to provide reasonable spatial reproducibility associations.

In addition to the previously described comparisons, we performed another validation of the model by considering an analysis where we use the WM task fMRI data and compare the 2-back working memory task periods to the 0-back working memory task ones. For each of these two different memory loading tasks there is a total of 330 time points of task ‘activation’ acquired, which according to Fig. 1, it is at the low side in temporal sampling, but it is sufficient given the HCP sample size to successfully use SPADE in this context. A common approach to this type of data is a traditional example of task fMRI data analysis where conditions are compared by means of a GLM analysis, and the results of such analyses in this same dataset have been previously reported in [22]. In this study, increased activation in the lateral-prefrontal and dorsal parietal cortex was associated with 2-back vs. 0-back working memory task differences. Table 2 shows the classification of the 10-fold cross validation results we obtained for discrimination of the 0-back and the 2-back working memory tasks. In all cases the classification performance was significantly better than random (permutation p<10^-3^). We appreciate that a twodimensional SPADE projection already provides a very high accuracy to identify which level of memory loading the task requires (2^nd^ column) at the single subject level. Such low dimensional representation already provides higher discrimination than the obtained using an SVM in full covariance space (5^th^ column). Further, in this scenario the two main filters provide a discrimination accuracy very close to the one obtained using all selected discriminant filters (3^rd^ and 4^th^ columns).

**Table 2:**
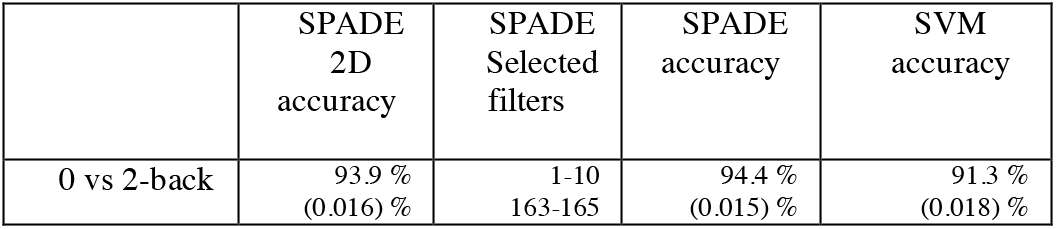
Summary results of the SPADE analyses in 2-back vs. 0-back working memory. The second column (SPADE 2-D) presents the classification results when discriminating each pair of fMRI modalities using a SPADE two-dimensional projection of the data. The column ‘SPADE selected filters’ shows the eigen-spectrum location of the filters selected by the model order selection. The column ‘SPADE accuracy’ shows the classification results obtained using the model order selection and the column ‘SVM accuracy’ shows the results obtained using an SVM in full covariance space (i.e. feature space of dimension (165*166)/2= 13695).

In Fig. 6 we present a summary of the spatial maps learned using the SPADE model. The filter providing the most variance for the 2-back task, i.e. filter 1, involves left thalamus and hippocampus, bilateral putamen and caudate, temporal pole and language areas. The next filter providing most variance for the 2-back task, i.e. filter 2, involves left thalamus, bilateral hippocampus, brainstem, cingulate and paracingulate gyrus, bilateral inferior frontal gyrus and left inferior temporal gyrus. On the other side of the eigen-spectrum, the filter maximizing variance for the 0-back task, i.e. filter 165, involves brainstem, cingulate gyrus, left thalamus, frontal pole and temporal gyrus and the next strongest filter for the 0-back task, i.e. filter 164, includes bilateral thalamus, brainstem, cingulate gyrus, frontal pole and motor cortex. Summarizing, the networks identified for the 2-back task involve areas purely related to working memory task while the 0-back task involves areas more related to the attention and salience network. This clearly reflects that the 2-back task involves stronger memory processing than the 0-back case which involves sustained attention. In general these presented results show some overlap with [23]. More specifically, some areas of filters 1 and 2 are present in both analyses. However, SPADE also reveals additional discriminative areas not found by the direct univariate analysis, including for example the striatum or the superior frontal gyrus.

**Fig. 6:**
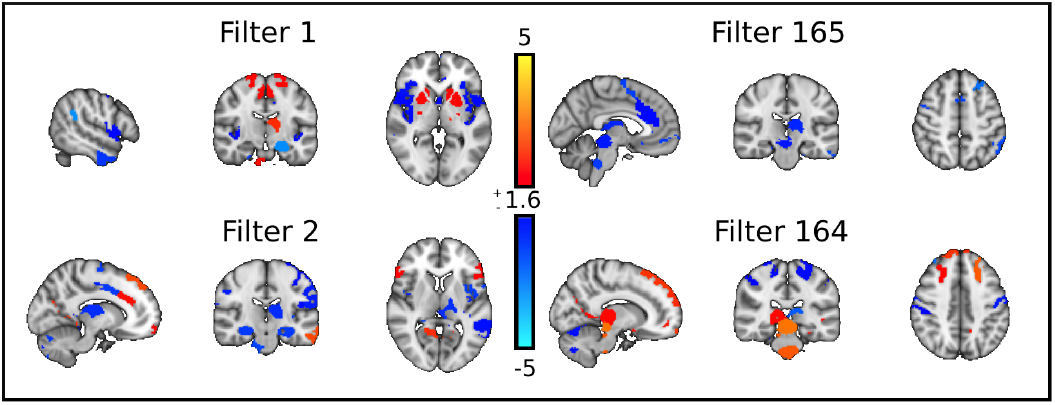
Spatial maps obtained using SPADE to compare the 2-back against the 0-back working memory fMRI data. Left side: filters 1 and 2 are the filters maximizing variance for the 2-back memory task. Right side: filters 164 and 165 are the ones maximizing variance for 0-back memory task.

## IV. Discussion

In this work we optimize and provide the tools to achieve optimal supervised linear filtering of fMRI data using the simultaneous diagonalization of covariance matrices [5], [6].

Although performing full brain voxel-wise analyses would be optimal, the high spatial resolution and correlation of fMRI data makes it computationally and technically challenging, and at the current state of development, an initial spatial dimensionality reduction is still required. Although different initial spatial dimensionality reductions are possible, including a parcellated full brain [14], [30], a resting state network space [3] or voxel-wise restricted to an anatomical ROI, here we use a full brain functional atlas parcellation [14] for ease of spatial validation of the results. We use regularized versions of the covariance matrices [15] and introduce a novel permutation strategy for model order estimation, i.e. how many spatial filters to select. Considering the estimated model order, the discriminatory generalization of the model is evaluated using cross-validation, it is tested for significance using permutation testing, and the associated interpretable spatial maps are obtained using structural coefficients [13]. We denote the fully developed pipeline as Spatial Patterns for Discriminative Estimation (SPADE, https://github.com/allera/SPADE).

We validated the use of this automated pipeline by using the high-quality HCP data from 1063 subjects. The analyses showed SPADE providing close to perfect low dimensional discrimination between representative fMRI experiments, and highlighted the straightforward interpretability of the model, with the obtained linear maps relating to connectivity between anatomical areas well known to be involved on each considered fMRI task. We also showed that accurate classification can be achieved also using (for example) an SVM in a highdimensional space (full covariance space).

However, the power of the presented model is the intuitive interpretation of the results that are summarized as a small set of spatial maps, each relating to a discriminative brain ‘sub-networks’, that can be used to further develop our understanding of brain function, directly or through its introduction in posterior analyses. This is certainly a big advantage with respect to discrimination in full covariance/correlation/partial correlation space where the interacting factors between different spatial nodes make the interpretation more difficult. As a note of advice, SPADE is not intended to replace classifiers on covariance-like feature spaces extracted from fMRI time-series; most probably, when not being able to achieve significant classification performance on covariance space when using some adequate and properly optimized classifier, SPADE will also not achieve significant discrimination. However, our experience suggests that if the classifier performs properly, then we probably will be able to learn more from the SPADE model than from the high dimensional classifier approach.

Our analyses revealed further a strong specificity of the spatial distribution across the three considered fMRI modalities (rest, motor and working memory). Each of the discriminative spatial maps obtained through SPADE reflects brain areas well-known to be relevant for a particular modality, reflecting for example that a stronger engagement of the DMN is recorded during rest than during task fMRI. In the working memory tasks, filters reveal independent effects for the ventral-dorsal pathways; the sensory integration is represented in one filter and the higher-order memory storage and utilization in another filter. Indeed, these two pathways start by sending sensory information to the parietal and temporal areas, characterizing spatial versus categorical features related to the items to memorize. Both connect to the <u>temporo-medial_structure</u> (TMS) via specific connectivity to the parahippocampus, enthorinal cortex, before converging on the hippocampus [31]. The TMS then connects through the temporal pole to the (prefrontal, pole) frontal and (anterior) cingulate to store and use the information. Regarding working memory, we observe the phonological loop with both Wernicke and Broca areas [32]–[34]. SPADE is extracting differentiating filters out of these networks, demonstrating the existence of independent dynamics that can be the subject of further studies.

We also presented extended analyses relating the discriminative generalization and the robustness of the learned spatial representations to the sample size and the fMRI temporal sampling. Although relatively high classification accuracy could be achieved using around 100 subjects, getting strongly reproducible all the significant spatial maps, required more than 200 subjects, probably even closer to 500 subjects. However, more in line with the classification results, the reproducibility of the four most discriminative maps could be achieved already with 100 subjects. Further, we observed that the top side of the eigen-spectrum achieves higher spatial reproducibility with less data than the bottom side. Although such behavior was not observed in all cases, it could be partially induced through the non-symmetric roles of the group covariances in the simultaneous diagonalization algorithm, and alternatives to further improve the spatial reproducibility in this direction are focus of current research.

To evaluate the performance of SPADE in a harder problem, we applied it to compare 2-back working memory from 0-back working memory task fMRI data. We were able to find the memory load of a subject with an accuracy higher than 94%. Further, the model showed that the 2-back task involves areas well known to be involved in working memory while the 0-back did not involve for example the hippocampus, but instead, the identified networks involved areas related to the salient and attention networks, as the frontal pole or the dorsal anterior cingulate gyrus. In comparing our results with those presented in [22], we note that our analysis point to connectivity changes between 2 and 0-back task at areas known to be involved in working memory tasks that were not clearly identified using the GLM approach in [22]. This is not a surprising result since both analyses are looking at different statistics in the data; while [22] reports changes in the mean fMRI signal across conditions through a GLM analyses, the findings we report using SPADE refer to changes in covariation between brain areas. Consequently, although the results are different, the analysis we present here and that in [22] are not exclusive but rather complementary, with both having their advantages and limitations. One of our research directions considers the development of SPADE towards single subject statistical spatial maps and voxel-wise representations, that could allow in the future its use to complement traditional GLM analyses at the first and second level.

Further, since the simultaneous diagonalization introduced by [5] can be used for any pair of symmetric positive definite matrices, and given the growing neuroimaging interest in working with partial correlation matrices, we have also explored the extension of the SPADE algorithm to partial correlation matrices and observe that it did not improve in discrimination with respect to the original formulation.

Nevertheless, we believe that such an approach warrants more thorough investigation in cases where the initial spatial dimensionality reduction results in more highly correlated data, for example when using a small spatially continuous ROI. The model is also very flexible in that it allows its implementation for different goals than direct discrimination, and it can for example be used to remove site-related variance from multi-site fMRI cohorts. Other research directions we are investigating for this model include the study of subject-wise deviation from group functional connectivity as a strategy to address normative modelling [35] on discriminative functional connectomes, as well as extensions to multi-class versions [36]. As an alternative to SPADE one could consider supervised non-linear dimensionality reduction models as the popular t-SNE [37], however, such models come with a loss of interpretation price.

To conclude, the SPADE model is able to capture and summarize the brain nodes involved in network covariation changes induced by fMRI experimental manipulation. As introduced here, the SPADE model can be used to further develop robust tools as well as to address questions of potential interest to diverse scientific and industrial communities.

## Funding

The research leading to the presented work has received funding from the developing Human Connectome Project (dHCP) through a Synergy Grant by the European Research Council under the European Union’s Seventh Framework Programme (FP/2007-2013), ERC Grant Agreement no. 319456. We further gratefully acknowledge support from the Netherlands Organization for Scientific Research (NWO) through VENI grant to J.N. (V1.Veni.194.032) and through VIDI grant to CFB (864.12.003). We also gratefully acknowledge funding from the Wellcome Trust UK Strategic Award (098369/Z/12/Z).

## Data availability and Ethics

Human Connectome Project data were acquired using protocols approved by the Washington University institutional review board and informed consent was obtained from all subjects. Anonymized data are publicly available from ConnectomeDB (db.humanconnectome.org). Informed consent and consent to publish was obtained from the Human Connectome Project according to the declaration of Helsinki. Research conducted at the Donders Center for Cognitive Neuroimage is covered by the protocol approved by the ‘Commissie Mensgebonden Onderzoek (CMO) Regio Arnhem-Nijmegen’ registered under CMO number 2014/288.

## Appendix

## A. Discriminative covariances diagonalization

Consider two sets of time-series, *X* and *Y*, measured at the same spatial locations during two different states of the system being measured, and define the spatial covariance matrices of each of these series as *C_x_* and *C_y_*. Usually, the unsupervised simultaneous diagonalization of both covariances is achieved through a generalized eigenvalue decomposition, involving a whitening transformation of one of the covariance structures (e.g. *C_x_*), combined with a further rotation to also diagonalize the other covariance (*C_y_*). A different approach was introduced in [5] where the whitening transformation is performed with respect to the sum of both covariances *C*:= *C_x_* + *C_y_*; such whitening can be summarized by two matrices, a rotation *W* and a scaling diagonal matrix *P*, such that *I* = *P^T^W^T^ CWP*. Individually applying these transformations to *C_x_* and *C_y_*, one obtains two (non-diagonal) matrices *K_x_* = *P^T^W^T^C_X_WP* and *K_Y_* = *P^T^W^T^C_Y_WP*. Performing now the eigenvalue decomposition of (for example) *K_Y_* we obtain matrices *Z_Y_* and diagonal *D_Y_* such that 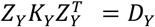. Defining *V* = *WPZ_Y_*, we have that *V^T^C_X_V* = *I* − *D_Y_* and *V^T^C_Y_V* = *D_Y_*. This means that *V* defines a basis for the data that diagonalizes simultaneously *C_X_* and *C_Y_* and, and by construction, the sum of the variances of the projections of data *X* and *Y* into each of the new basis vectors adds to one; consequently, for basis directions where the projection of *X* has large variance (i.e. close to one), the projection of *Y* must have low variance (i.e. close to zero), and vice versa. The algorithmic imposition to maximize variance for one class and minimize it for the other one, makes of this process a supervised multivariate dimensionality reduction model, and the new learned basis is formed of spatial filters that provide optimal linear discrimination in terms of variance of the projected data [12]. Note that for implementation purposes it all reduces to solving a generalized eigenvalue problem which can be easily accomplished using most software platforms, for example using the matlab command *eig*(*C_X_, C*) or the python command *scipy. linalg. eigh*(*C_X_, C, eigvals_only* = *False*). In the recent literature, the process described in [5] is commonly used as a feature extractor for EEG based brain computer interfaces [11], [12] and the features extracted for discrimination are usually the logarithmic of the variance of the projected data; by construction of the algorithm the most discriminative dimensions are the associated with the extreme eigenvectors since towards the center of the eigen-spectrum the variances for both conditions tend to be close to 1/2. In practice, for discrimination purposes the dimension is usually reduced to a few pairs of extreme eigenvectors and a low dimensional linear classifier suffices [12].

## B. Interpretation of the linear filters

Since the simultaneous diagonalization assumes no implicit noise model, the spatial filters learned, i.e. the columns of *V* (*V_k_*) cannot be directly interpreted and the interpretable spatial maps (*A_K_*) are obtained from *A* = *C_X|Y_VC_S_*^−1^ [13], [18], where *X|Y* denotes the spatially concatenated *X* and *Y* data, and *S* the data projected to the new basis *V*, i.e. *S* = *V^T^*(*X|Y*). For completeness we note here that *X|Y* = *AS* + *e*, where *e* denotes the residual of the approximation *X|Y* = *AS* and for a more detailed description we refer the reader to [13], [18].

## C. Model order estimation

Although for classification purposes the SPADE model order selection is not critical due to the latter integration of a classifier, for interpretation purposes we are required to select which filters provide significant connectivity covariation information. To that end we address for the first time the model order estimation on this model by permutation testing to find a robust estimation of which filters explain more variance for a modality than for the other one, with respect to a null distribution obtained by permutation.

To that end a p-value is computed for each basis vector (column of *V, V_k_*) by comparing the absolute difference between the variances of the class-wise data projections, 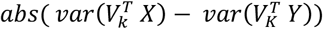, with a distribution of such values obtained when the simultaneous diagonalization basis (V) is computed using randomized groups of covariances. In the dataset set used in the main text the randomization is performed 1000 times and it is achieved by permuting half of the subject’s covariance matrices across different fMRI modalities. We consider a filter significant at a Bonferroni corrected level p<0.05/165.

## D. On the spatial reproducibility

Fig. A1 shows the standard deviation of the correlation matrices values presented in Fig. 4. We observe that the standard deviation of all elements in general, and of the diagonal elements in particular decreases with a bigger sample size.

**Fig. A1:**
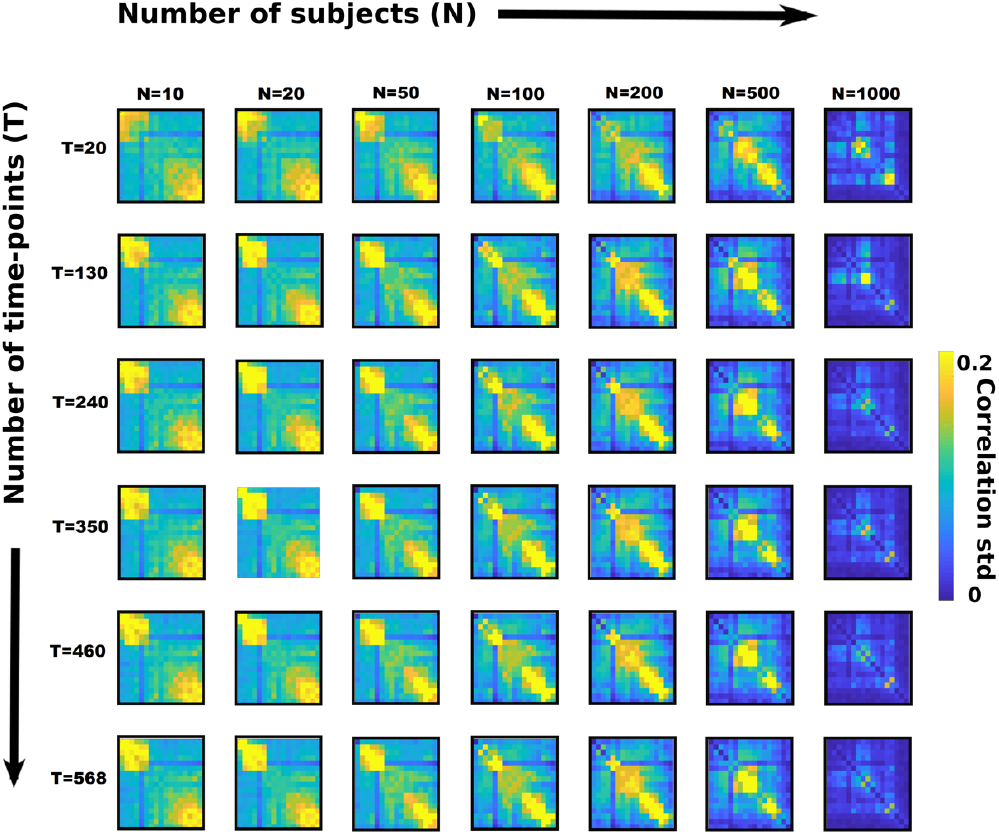
Standard deviation of SPADE spatial maps reproducibility for RT vs MT. Each subfigure shows the element wise standard deviation around the correspondent mean correlation shown in Fig. 4. The standard deviation is computed over 1000 bootstraps (see Methods section for details).

## Acknowledgment

Data were provided by the Human Connectome Project, WU-Minn Consortium (Principal Investigators: David Van Essen and Kamil Ugurbil; 1U54MH091657) funded by the 16 NIH Institutes and Centers that support the NIH Blueprint for Neuroscience Research; and by the McDonnell Center for Systems Neuroscience at Washington University.

## Notes

The authors declare to have no conflict of interests.

### Competing Interest Statement

The authors have declared no competing interest.

### Summary of Updates

Add experiments to study spatial reproducibility of results

